# CDC6 regulates both G2/M transition and metaphase-to-anaphase transition during the first meiosis of mouse oocytes

**DOI:** 10.1101/587238

**Authors:** Zi-Yun Yi, Tie-Gang Meng, Xue-Shan Ma, Jian Li, Chun-Hui Zhang, Ying-Chun Ouyang, Heide Schatten, Jie Qiao, Qing-Yuan Sun, Wei-Ping Qian

**Affiliations:** The Reproductive Medicine Center; Peking University Shenzhen Hospital; Shenzhen Peking University-The Hong Kong University of Science and Technology Medical Center; Shenzhen, Guangdong 518036, China; State Key Laboratory of Stem Cell and Reproductive Biology; Institute of Zoology; Chinese Academy of Sciences; Beijing 100101, China; Department of Veterinary Pathobiology; University of Missouri; Columbia, MO 65211, USA; Reproductive Medical Center, Peking University Third Hospital, Beijing 100191, China

**Keywords:** CDC6, oocyte, meiosis, germinal vesicle breakdown, spindle

## Abstract

Cell division cycle protein CDC6 is essential for the initiation of DNA replication. CDC6 was recently shown to inhibit the microtubule-organizing activity of the centrosome. Here, we show that CDC6 is localized to the spindle from Pro-MI to MII stages of oocytes, and it plays important roles at two critical steps of oocyte meiotic maturation. CDC6 depletion facilitated the G2/M transition (GV breakdown, GVBD) through regulation of Cdh1 and cyclin B1 expression and CDK1 phosphorylation in a GVBD-inhibiting culture system containing milrinone. Furthermore, GVBD was significantly decreased after knockdown of cyclin B1 in CDC6-depleted oocytes, indicating that the effect of CDC6 loss on GVBD stimulation was mediated, at least in part, by raising cyclin B1. Knockdown of CDC6 also caused abnormal localization of γ-tubulin, resulting in defective spindles, misaligned chromosomes, cyclin B1 accumulation and spindle assembly checkpoint (SAC) activation, leading to significant Pro-MI/MI arrest and PB1 extrusion failure. These phenotypes were also confirmed by time-lapse live cell imaging analysis. The results indicate that CDC6 is indispensable for maintaining G2 arrest of meiosis and functions in G2/M checkpoint regulation in mouse oocytes. Moreover, CDC6 is also a key player regulating meiotic spindle assembly and metaphase-to-anaphase transition in meiotic oocytes.

**Summary statement:** We show that CDC6 is indispensable for maintaining G2 arrest of mouse oocytes. Moreover, CDC6 is also a key player regulating meiotic spindle assembly and metaphase-to-anaphase transition in meiotic oocytes.

## Introduction

Mammalian oocytes are arrested at the first meiotic prophase, also termed the germinal vesicle (GV) stage. Germinal vesicle breakdown (GVBD) marks the initiation of oocyte maturation, followed by microtubule assembly around chromosomes during pro-metaphase I, chromosome migration to the central plate of the spindle during the MI stage, and subsequent metaphase-to-anaphase transition and final extrusion of the first polar body (PB1). The prophase I arrest and progression from metaphase I (MI) to metaphase II (MII) are two critical stages during oocyte maturation (Hayden et al., 2009).

Resumption of meiosis, referred to as GVBD is mainly associated with regulation of the maturation-promoting factor (MPF) activity, which is a complex of regulatory subunit cyclin B1 (CCNB1) and the catalytic subunit cyclin-dependent –kinase 1 (CDK1), also known as Cell Division Cycle 2 (CDC2) (Heikinheimo and Gibbons, 1998; Adhikari et al., 2012). Activation of CDK1 takes place through the association with cyclins, suppression of CDK1-inhibiting kinases (Myt1/Wee1) and activation of CDK1-activating phosphatases (CDC25) (David, 1995; Han et al., 2006). Wee1/Myt1 kinase family inhibits CDK1 activity by phosphorylating its Thr14 and Tyr15 residues (Morgan, 1995; Jones, 2011). A high level of cAMP activates protein kinase A (PKA), which is likely to maintain prophase I arrest by activating Wee1 B (Han et al., 2005). CDK1 can also be activated by the Cdc25 phosphatases, including Cdc25A, cdc25B, and Cdc25C in mammals, dephosphorylating Wee1 B-phosphorylated CDK1 (Branzei and Foiani, 2008; Lincoln et al., 2002). The G2 arrest in meiosis is dependent not only on CDK1 activity, but also on regulating concentrations of cyclin B1. Cyclin B1 degradation during prophase I arrest is induced by Cdh1-mediated activation of the APC (Holt et al., 2011, Reis et al., 2006). Meiotic competency of GV oocytes was accelerated by Cdh1 loss (Holt et al., 2011). Emil and securin modulators of APC/C^cdh1^ activity can inhibit cyclin B1 degradation and improve activation of MPF (Marangos et al., 2007). Interestingly, the spindle checkpoint protein BubR1 has also been shown to regulate prophase I arrest. Knockdown of BubR1 in GV oocyte can release the oocytes from prophase I arrest in the presence of 3-isobutyl-2-methy I xanthine (IBMX) and arrest the oocytes at the pro-metaphase I stage by decreasing the expression level of Cdh1 (FZR1), which is a well-established co-activator of APC/C (Hayden et al., 2009).

CDC6 (cell division cycle-6 homologue), was first identified as an important protein for the initiation of DNA replication. It is indispensable for the formation of pre-RCs (pre-replication complexes) (Coleman, 2002, Rowles et al., 1999). Different strategies regulate accurately the stability and cellular location of the CDC6 protein (Piatti et al., 1996, Sánchez et al., 1999). In human cells, phosphorylation of CDC6 by cyclin A-CDKs leads to a down-regulation of CDC6 activity during the S phase by translocation from the nucleus into the cytoplasm (Petersen et al., 2014). In the budding yeast, the mitotic CDK and cyclin B bind specifically to the amino-terminal domain (NTD) of CDC6, and CDC6 in the complex is unable to assemble pre-RCs (Mimura et al., 2004). In *Xenopus* oocytes, during transition from MI to MII, CDC6 is under the antagonistic regulation of cyclin B (which interacts with and stabilizes CDC6) and the Mos-MAPK pathway (which negatively controls CDC6 accumulation) (Daldello et al., 2015).

Once DNA is licensed to undergo replication, CDC6 is no longer required for the continued association of MCM with DNA and is removed from the pre-RCs. Except for its function during DNA synthesis, CDC6 also plays a role later in the cell cycle in mitosis. Overexpression of CDC6 during the late S phase leads to mitotic arrest at the G2/M-phase transition, by preventing cyclin B/CDK1 activation (Lorena et al., 2014). CDC6 also participates in the regulation of exit from mitosis, by stabilizing the anaphase-promoting complex/cyclosome (APC/C) substrate and affecting modulation of APC/C^CDC20^ (Boronat and Campbell, 2007). In Hela cells, CDC6 and Plk1 colocalize at the spindle pole in metaphase and at the central spindle in anaphase. Furthermore Plk1-mediated phosphorylation of CDC6 is required for CDK1 inhibition, release of separase, and subsequent anaphase progression (Hyungshin and Erikson, 2010). The mouse CDC6 binds chromatin and associates with the spindle during mitosis. A recent study suggests a novel function for Cdc6 in coordinating centrosome assembly and function (Lee et al., 2017).

These results indicate that CDC6 may have important roles in regulating cell cycle progression in addition to the initiation of eukaryotic DNA replication. We showed that depletion of CDC6 facilitated G2/M transition by elevating cyclin B1 levels and decreasing Cdh1 levels. While, the accelerated G2/M transition induced by CDC6 knockdown could be rescued by injecting cyclin B1 siRNA. Furthermore, our results showed that CDC6 localized along the meiotic spindle and plays a crucial role in spindle assembly, metaphase-to-anaphase transition and subsequent first polar body extrusion in meiotic oocytes.

## Results

### Expression and subcellular localization of CDC6 during oocyte meiotic maturation

We collected oocytes cultured for 0, 4, 8 and 12h, corresponding to germinal vesicle (GV), germinal vesicle breakdown (GVBD), metaphase I and metaphase II stages, respectively, to examine the expression level and subcellular localization of CDC6 during meiosis. The CDC6 mRNA level was measured by real-time PCR (RT-PCR). We found that, compared to the GV oocytes, the mRNA levels of CDC6 in GVBD, MI and MII oocytes were notably decreased (35.37±2.40%, 43.55±2.03%, 35.31± 3.39% VS 100%). As shown in Figure 1B, Western blot results showed that CDC6 was expressed in all stages at a similar level at the GV, GVBD and MI stages, while its expression decreased at the MII stage in mouse oocytes. To investigate the subcellular localization of CDC6 during meiotic maturation, oocytes in different stages were processed for immunofluorescent staining. As shown in Figure 1C, CDC6 mainly localized in the germinal vesicle at the GV stage. After GVBD, CDC6 gradually accumulated in the vicinity of condensed chromosomes. At MI and MII, CDC6 was found to distribute on spindle microtubules. These results suggest that CDC6 might function in mouse meiotic maturation.

**Figure 1:**
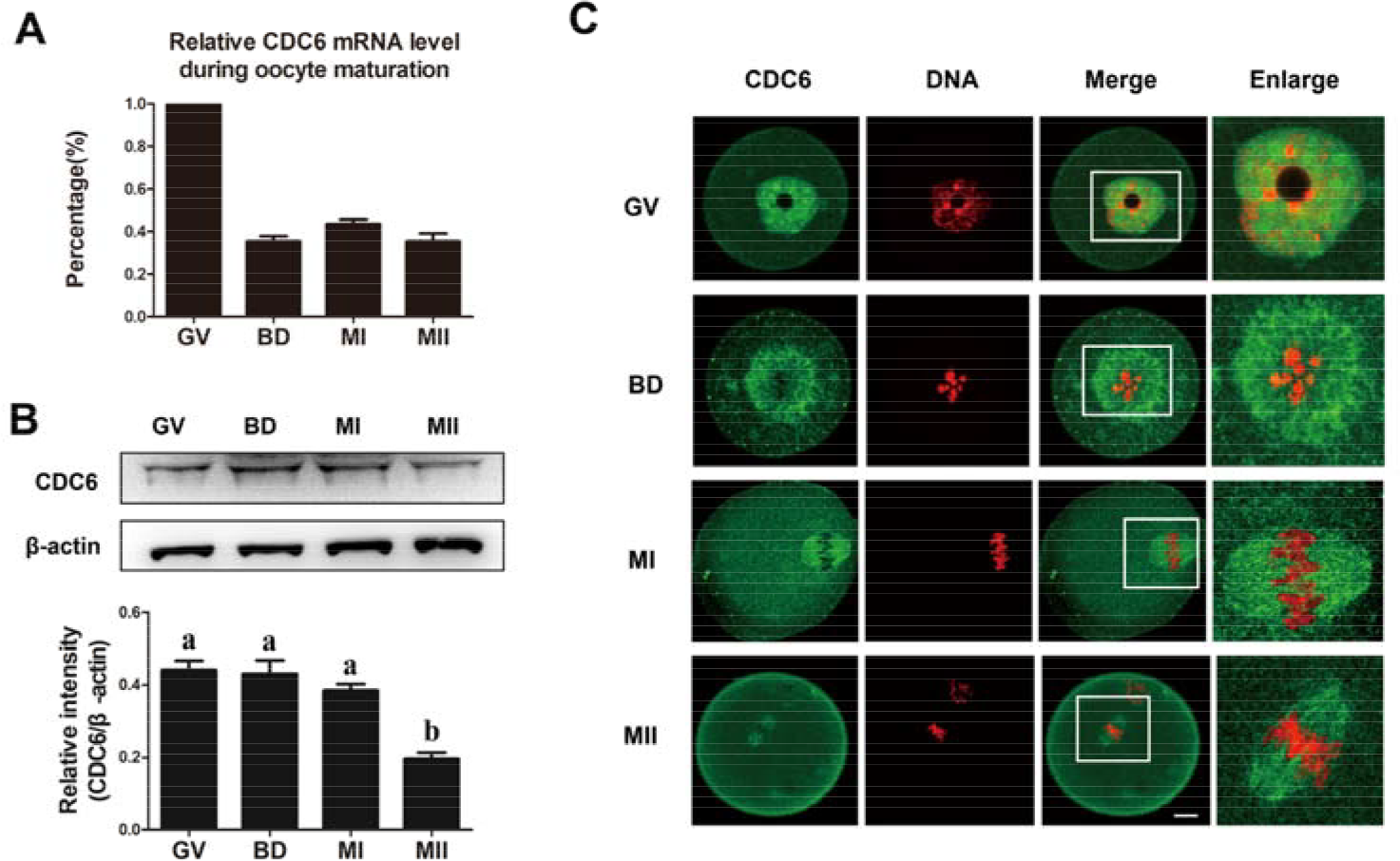
Expression and subcellular localization of CDC6 during meiotic maturation in mouse oocytes. (A) Relative level of CDC6 mRNA identified by RT-PCR. CDC6 mRNA levels were normalized to the maximum levels at the GV stage. A total of 50 oocytes were collected after culture for 0, 4, 8 and 12h, representing GV, GVBD, MI and MII stages, respectively. (B) Expression level of CDC6 was detected by Western blotting. Samples were collected after 0, 4, 8 or 12h of culture, corresponding to the GV, GVBD, MI and MII stages, respectively. The molecular weight of CDC6 and β-actin were about 63 kDa and 43 kDa, respectively. Each sample contained 200 oocytes. The relative level staining intensity of CDC6 was accessed by densitometry. The relative intensity of CDC6 was accessed by gray-scale analysis using the software Quantity One (Bio-rad). Levels of expression were normalized to the levels of β-actin. Error bars are mean± SEM. (C) Subcellular localization of CDC6 as revealed by immunofluorescent staining. Oocytes at the GV, GVBD, MI and MII stages were stained with antibody against-CDC6 (green). Oocytes were stained with Hoechst 33342 to visualize DNA (red). Bar=20μm.

### Depletion of CDC6 facilitates GVBD and CDK1 activity dependent on cyclin B1

To examine the role of CDC6 in mouse oocyte maturation, we knocked down CDC6 by its specific siRNA injection. Western blot revealed that CDC6 level was notably reduced in the CDC6 siRNA-injected group (0.12 ± 0.014) compared with the control siRNA injected group (0.51± 0.029, P<0.001, Fig.2A). Next, we stained the CDC6 siRNA-injected oocytes with antibody against CDC6, and there was little or no immunostaining in the germinal vesicle (Fig. 2B). The CDC6 siRNA-injected oocytes were arrested in M2 medium containing 0.75μM milrinone (which was the minimum concentration of milrinone to maintain meiotic arrest in mouse oocytes;(Duncan et al., 2009)), for up to 24h to count the number of GVBD oocytes at different time points. The percentage of oocytes at the GVBD stage was considerably higher in the CDC6 knockdown group than in control oocytes (6h GVBD rate, 24.26±2.65 % versus 6.12± 1.34%; 12h GVBD rate, 43.9±2.25% versus 15.86± 1.69 %; 18h GVBD rate, 65.33± 5.26% versus 21.82± 2.46%; 24h GVBD rate, 71.58± 4.21% versus 24.2± 2.91%; P<0.05; Fig.2C).

**Figure 2.**
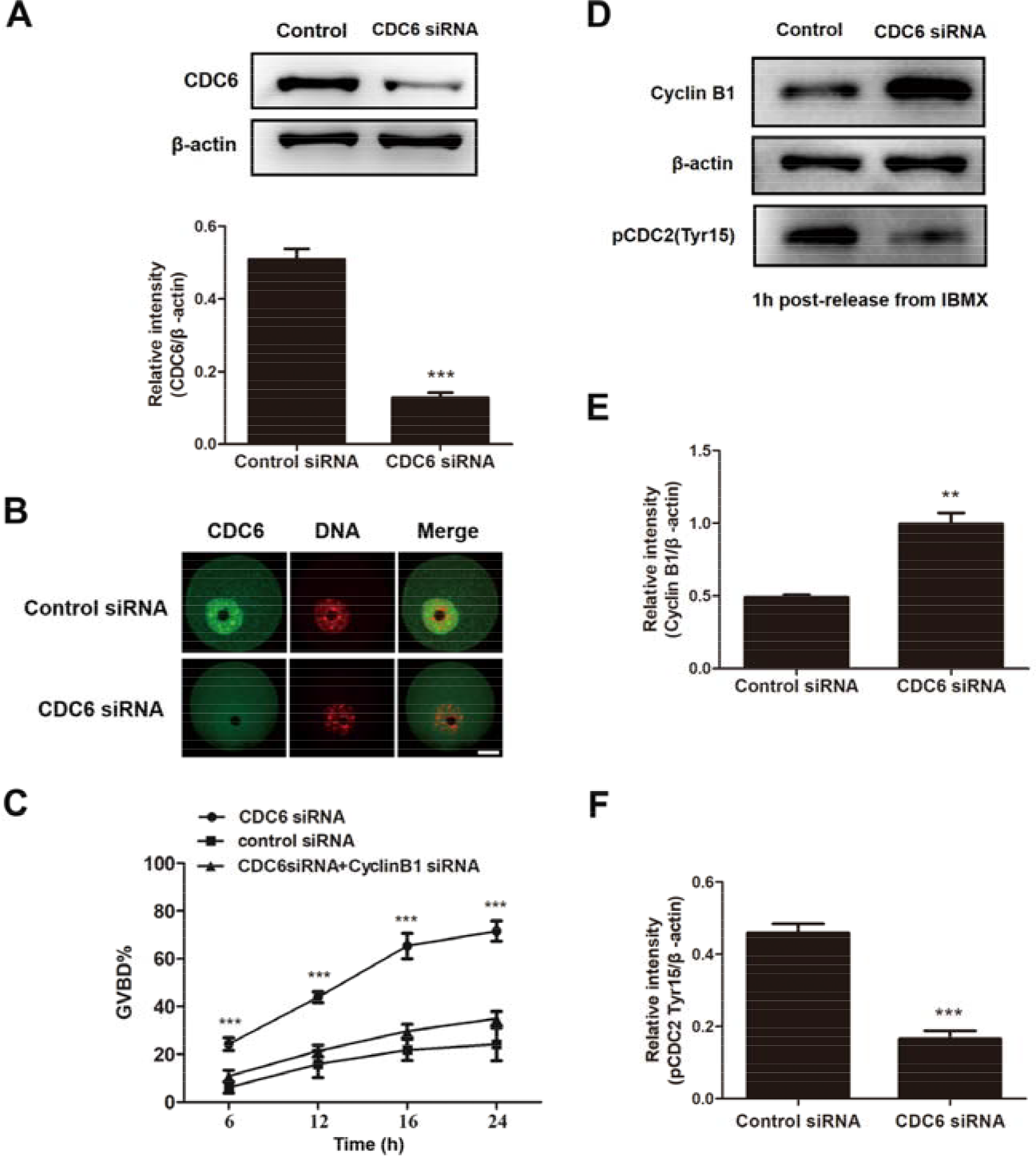
CDC6 depletion accelerated meiotic resumption and activated MPF activity in mouse oocytes. (A) Western blot of CDC6 in the oocytes microinjected with CDC6 siRNA or control siRNA; the oocytes were arrested in M2 medium containing 200μM IBMX for 24h. The molecular mass of CDC6 is 63kDa and that of β-actin is 43kDa. A total of 200 oocytes were used in each lane. The relative intensity of CDC6 was assessed by gray-scale analysis using the software Quantity One (Bio-Rad). Levels of staining intensity were normalized to levels of β-actin. Data are presented as mean±SEM of 3 independent experiments (***,p<0.001). (B) Confocal microscopy showing depletion of CDC6 protein after siRNA. A total of 52 oocytes were assessed in the CDC6 siRNA injected group and 46 oocytes were assessed in the control-siRNA injected group. (C) The GVBD rates at 6, 12, 18, 24h following arrest in M2 medium containing 0.75μM milrinone for the CDC6 siRNA-injected, control siRNA injected and CDC6 siRNA +cyclin B1 siRNA-injected group. Oocytes microinjected with CDC6 siRNA, control siRNA or CDC6 siRNA +cyclin B1 siRNA were arrested in M2 medium containing 200μM IBMX for 24h, then transferred to M2 medium containing 0.75μM milrinone. (D) Western blotting of CDC6, cyclinB1, pCDK1 (Tyr15 of CDK1) and β-actin in CDC6 siRNA and control siRNA injected oocytes, 1h after release from IBMX. A total of 200 oocytes were used in each sample. CDC6 is 63kDa, cyclinB1 is 55kDa, pCDK1 is 34kDa and β-actin is 43kDa. (E and F) Gray-scale analysis of cyclinB1 and pCDK1. The relative intensity of cyclin B1 or pCDK1 was assessed by gray-scale analysis using the software Quantity One (Bio-Rad). Levels of staining intensity were normalized to levels of β-actin. Data are presented as mean±SEM of 3 independent experiments (***,p<0.001).

Furthermore, we tested CDC2 activity by examining its Tyr15 phosphorylation state. The activity of CDC2, which was required for MPF activation, was higher in the CDC6-depleted oocytes (0.166 ± 0.022) compared to that of the control group (0.46 ± 0.025, P<0.05, Fig. 2D,F) by 1h following release from IBMX. We then determined whether CDC6 depletion would affect the expression of cyclin B1, which is indispensable for CDC2 and MPF activation. The level of cyclin B1 in the CDC6-depleted oocytes (0.99± 0.076) was notably higher than that in the control oocytes (0.49± 0.017, P<0.05, Fig. 2D, E). These data suggest that CDC6 is required for destabilizing cyclin B1 and inhibiting MPF activity, and is thus necessary to maintain prophase arrest.

### GVBD induced by CDC6 depletion can be rescued by cyclin B1 knockdown

We next determined whether cyclin B1 depletion could rescue the facilitated G2/M transition. We mixed CDC6 and cyclin B1 siRNAs and microinjected the mixture into the GV oocytes. This achieved a 70% depletion of cyclin B1 expression level (Fig. S1). We found that the GVBD rate of the co-RNAi group (24h GVBD rate, 34.99 **±** 2.98 %) was lower than that of the CDC6 siRNA injected oocytes (24h GVBD rate, 71.58 ± 4.21%; P<0.05; Fig.2C).

### Knockdown of CDC6 causes up-regulation of cyclin B1 level by inhibiting the APC/C^cdh1^-mediated destruction

To examine more directly if CDC6 depletion causes increased levels of cyclin B1, control and CDC6 depletion oocytes were microinjected with exogenous cyclin B1-GFP mRNA (20ng/μl) and cultured in 0.75μM milrinone for 10 hours, and then the dynamics of cyclin B1-GFP accumulation was assessed. There was a significant rise in the cyclin B1 level in CDC6-depleted oocytes compared with the control oocytes. (Fig. 3. A, B; P<0.05).Furthermore, the accelerated GVBD observed in CDC6-depleted oocytes can be ameliorated by cyclin B1 knockdown. Together, these results indicate that the effects of CDC6 knockdown in oocytes are mediated, at least in part, by elevated cyclin B1.

**Figure 3.**
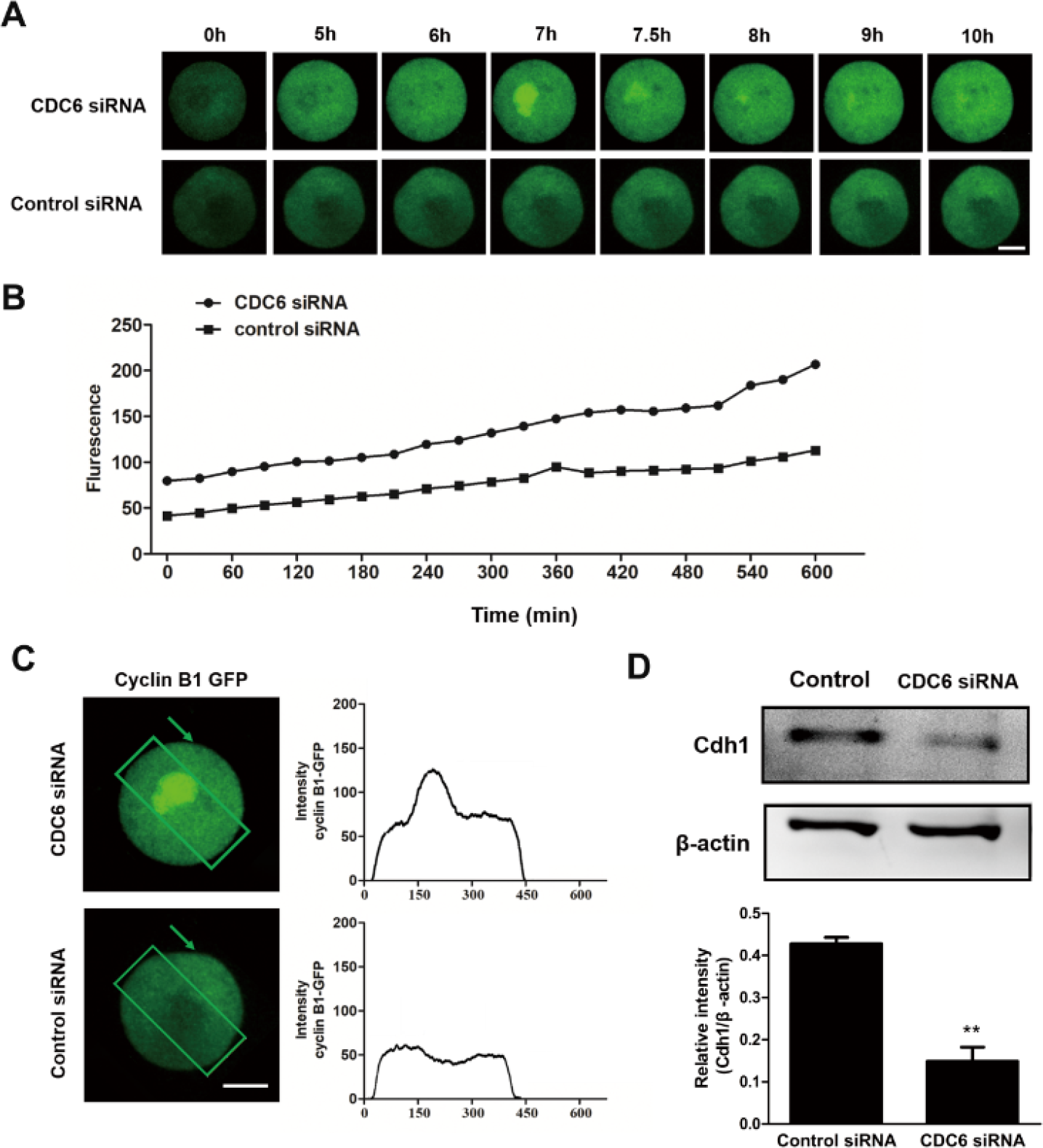
CDC6 knockdown causes increase of cyclin B1 level, accelerates nuclear translocation of cyclin B1 before GVBD and decreases level of Cdh1. First, CDC6 siRNA or control siRNA was injected into the GV oocytes. After a 24h incubation in 200μM IBMX, the oocytes were injected with cyclin B1-GFP mRNA and arrested for 15min in 200μM IBMX, followed by culture in M2 medium containing 0.75μM milrinone for 10h. Time points indicate the time lapse from about 15min after being injected with cyclin B1-GFP mRNA. The fluorescence intensity of cyclin B1-GFP at all time points were quantified by Image J (n=30 cells for each quantification). (A) Representative images of nuclear localization pattern of cyclin B1-GFP in CDC6 siRNA and control siRNA injected groups. (B) Time-dependent changes in cyclin B1-GFP fluorescence intensity. (C) The boxed area across the oocyte was assessed for fluorescence intensity by Image J. (D) Western blot of Cdh1 in the oocytes injected with CDC6 or control siRNA, 1h after release from IBMX. Cdh1 is 55kDa and β-actin is 43kDa. Gray-scale analysis of Cdh1/β-actin was assessed by software Quantity one (Bio-rad). Data are presented as mean ±SEM of 3 independent experiments. (**, p<0.01)

Elevated nuclear cyclin B1 levels, an event normally observed in oocytes before nuclear envelope breakdown, is a prerequisite for MPF activation and the occurrence of G2/M transition (Holt et al., 2010). Live cell imaging demonstrated the ability of the cyclin B1 GFP to accumulate in the nucleus in the CDC6-depleted oocytes. While, in the control group, cyclin B1-GFP failed to accumulate in the nucleus in the presence of 0.75μM milrinone (Fig. 3C).

In mammalian oocytes, APC/C^cdh1^ activity is required to repress cyclin B1 levels in oocytes during prophase I arrest, maintaining the G2/M arrest in meiosis (Holt et al., 2011). We found that the level of Cdh1 in the CDC6-depleted oocytes (0.149**±** 0.0331) was notably lower than that in the control oocytes (0.428**±** 0.014, P<0.05, Fig. 2D). Thus, it is likely that reduction of CDC6 can accelerate the G2/M transition by decreasing the expression level of Cdh1.

### Depletion of CDC6 causes abnormal spindles and misaligned chromosomes

To further clarify the roles of CDC6 in mouse oocyte meiotic maturation, we determined spindle morphology and chromosome distribution using immunofluorescent staining. In the CDC6 depletion group, oocytes exhibited various kinds of morphologically defective spindles and misaligned chromosomes. The major spindle defects were mini-spindle and spindles with abnormal poles, including spindles with no poles, one pole, and multiple poles (Fig.4A). Minor chromosome misalignments, the absence of chromosome alignment or lagging chromosomes were observed (Fig. 4A).

**Figure 4.**
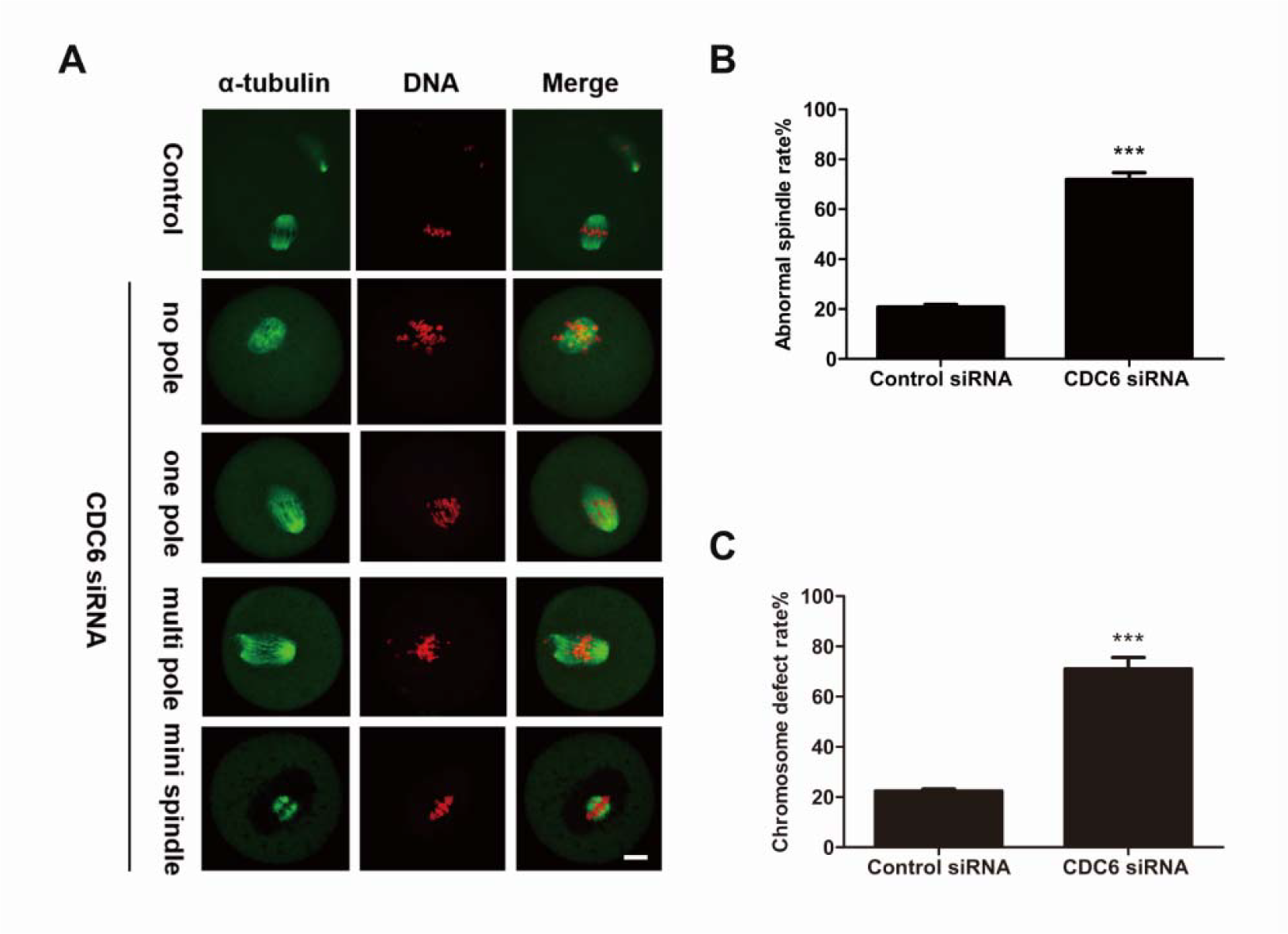
Depletion of CDC6 causes severely abnormal spindle and misaligned chromosomes in mouse oocytes. (A) Oocytes microinjected with CDC6 or control siRNA were incubated in M2 medium containing 200μM IBMX for 24h, then washed and transferred to IBMX-free M2 for 14h, followed by immunostaining with α-tubulin (green) and Hoechst 33342 (red). CDC6-depleted oocytes exhibited various morphologically defective spindles and misaligned chromosomes. Bar=20μm. (B and C) Percentage of oocytes with abnormal spindles and misaligned chromosomes in the CDC6 depletion group and control group. Data are presented as mean±SEM of 3 independent experiments (***,p<0.001).

As shown in Figure 4B, the proportion of oocytes with abnormal spindles in the CDC6-siRNA injection group (71.86 ± 2.73 %,n=124) was significantly higher than that in the control-siRNA injection group (20.79 **±**1.678 %,n=116; P<0.001). Moreover, as shown in Figure 4C, the rate of misaligned chromosomes in the CDC6 depletion group (71.06 **±**4.498%, n=124) and control groups (22.39 **±**0.289 %, n=116) differed significantly (P<0.01).

### Knockdown of CDC6 leads to dissociation of γ-tubulin from spindle poles

The above results revealed that CDC6 might be involved in spindle organization and spindle pole congregation. It is well known that the MTOC-associated protein γ-tubulin is a key regulator in microtubule nucleation and organization in acentriolar mouse oocytes. Since CDC6 partially colocalizes with γ-tubulin, we further investigated the effect of CDC6 knockdown on γ-tubulin localization. As shown in Figure 5, γ-tubulin was localized to the spindle poles in the control siRNA-injected oocytes (n=75) at the MI stage. In contrast, the depletion of CDC6 caused dissociation of γ-tubulin form the disrupted spindle poles, while it became irregularly distributed at the spindle fibers or dispersed into the cytoplasm (n=67).

**Figure 5.**
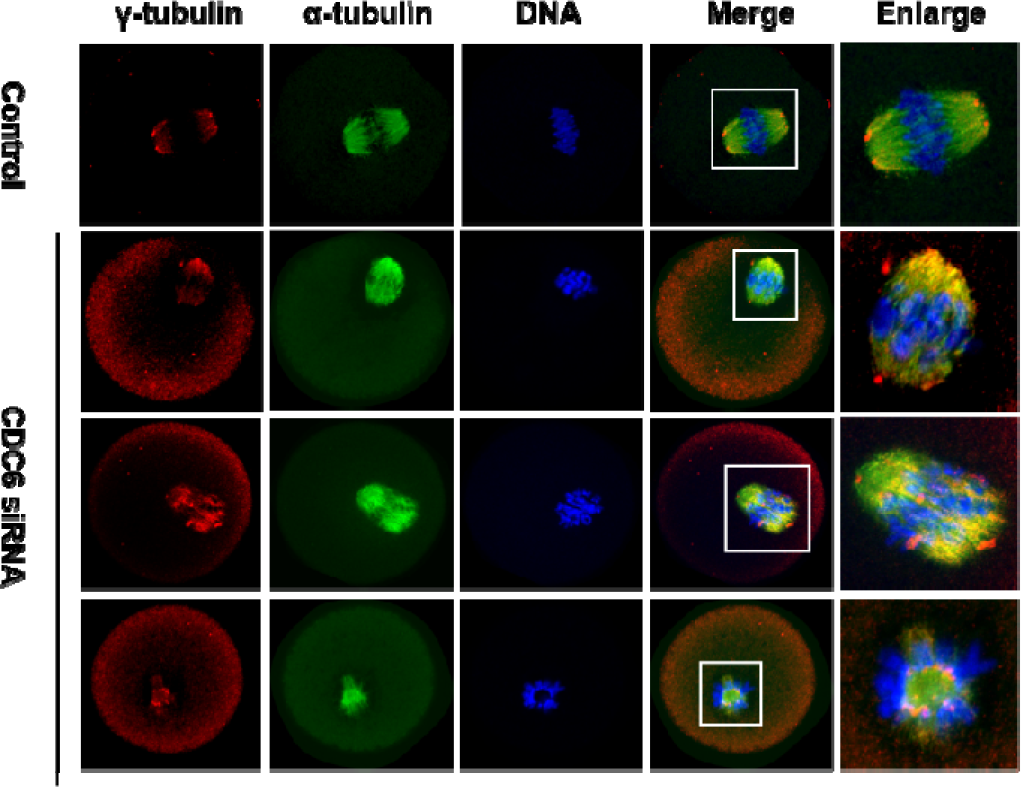
Dislocation of γ-tubulin from spindle poles in CDC6-depleted oocytes. Oocytes injected with CDC6 siRNA or control siRNA were incubated in fresh M2 medium for 8h after being arrested in M2 medium containing 200μM IBMX for 24h, followed by staining of α-tubulin (green), γ-tubulin (red) and DNA (blue). In the control siRNA-injected group, γ-tubulin was associated with the spindle poles at the MI stage, whereas γ-tubulin dissociated from the abnormal spindle poles and dispersed into the cytoplasm in the CDC6-siRNA injected group. α-tubulin, green; γ-tubulin, red; DNA, blue. Bar=20μm.

### CDC6 knockdown causes increased levels of cyclin B1, pro-MI/MI arrest, and reduced PB1 extrusion

After microinjection of CDC6 siRNA or control siRNA, oocytes were maintained in M2 medium containing 200μM IBMX for 24h; then oocytes were continuously cultured in IBMX-free M2 medium for 10 or 14h. After culture for 10h, we found that the majority of CDC6 depleted oocytes were arrested at the Pro-MI/MI stage, while most oocytes in the control-siRNA group passed through the MI stage (Fig, 6A). The pro-MI/MI arrest rate in the CDC6-depletion group (72.79 **±** 2.108%, n=94) was considerably higher than that in the control group (34.76 ± 1.508 %, n=97, P<0.01, Fig. 6B).

**Figure 6.**
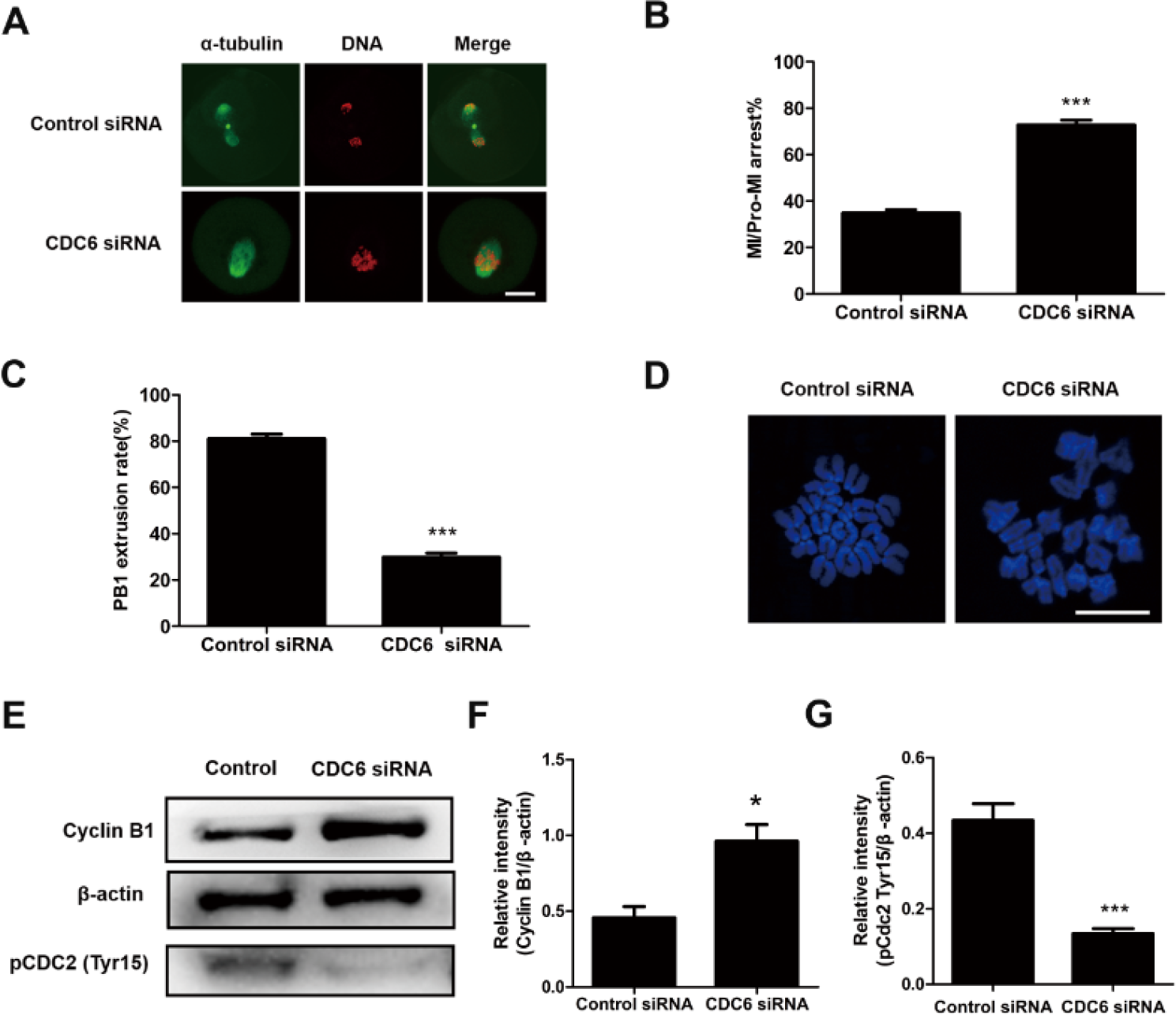
CDC6 depletion causes Pro-MI/MI arrest, increase of cyclin B1 level and failure of PB1 extrusion. After microinjection of CDC6 siRNA or control siRNA, the oocytes were incubated in M2 medium containing IBM for 24h, then washed and transferred to IBMX-free M2 medium for 10 or 14h. (A) Oocytes microinjected with CDC6 or control siRNA were cultured for 10h, followed by immunostaining with α-tubulin and staining with Hoechst 33342. In the CDC6 depletion group, oocytes were arrested at the Pro-MI/MI stage, but oocytes in the control group had reached the T1 (A1:Syt1) stage. α-tubulin, green; DNA, red. Bar=20μm. (B) Rates of Pro-MI/MI cultured oocytes, when CDC6 siRNA or control siRNA-injected oocytes were cultured for 10h after release from IBMX. Data are presented as mean±SEM of at least three independent experiments (***, p<0.001). (C) Percentage of first polar body extrusion in the CDC6 knockdown group and control group. Data are presented as mean±SEM of 3 independent experiments (***, p<0.001). (D) Oocytes of CDC6 siRNA and control siRNA injected group were cultured for 12h, followed by chromosome spreads. Bar=20μm. (E) Western blot showing expression levels of cyclin B1 and pCDC2 (Tyr15 of CDC2) in the CDC6 knockdown group and control group. Each sample contained 150 oocytes. (F and G) The intensity of cyclin B1/β-actin and pCDC2/β-actin were assessed by grey level analysis using the software Quantity one (Bio-Rad). Data are presented as mean±SEM of 3 independent experiments (*,p<0.05) (***,p<0.001).

As shown in Figure 6C, after culture for 14h, the rate of PB1 extrusion in the CDC6 -knockdown group (81.66±1.945 %, n=115) was remarkably lower than that in the control group (29.9±1.71 %, n= 127, P<0.01, Fig.5C). Since the majority of CDC6-depleted oocytes were arrested at the pro-MI /MI stage with abnormal spindles and misaligned chromosomes (Fig.,4 A) we asked whether chromosomes could segregate correctly. As shown in Figure 6D, chromosome spreading test indicated that chromosomes were still in the bivalent state in the CDC6-depleted oocytes even after 14h of culture; while univalent chromosomes could be observed in the control oocytes, suggesting completion of anaphase.

To further explore the reason for the pro-MI/MI arrest in CDC6 depleted oocytes, the level of cyclin B1 was examined at 10h of culture in IBMX-free M2 medium. As shown in Figure 6E, the cyclin B1 expression level was significantly higher in the CDC6 depletion group compared to the control group, indicating a failure of degradation of cyclin B1 in the CDC6 knockdown group, and thus Pro-MI/MI arrest. Once oocytes reach MI, the degradation of cyclin B1 inactivates of CDK1, which can promote anaphase I and exit from meiosis I. Furthermore, we tested CDK1 activity by examining its Tyr15 phosphorylation state. The activity of CDK1, was higher in the CDC6-depleted oocytes.

### CDC6 depletion causes activation of the SAC

The CDC6 knockdown oocytes failed to undergo correct homologous chromosome segregation, which prompted us to analyze the localization of the SAC protein Bub3 and Mad1. Specific signals for Bub3 were observed in CDC6 knockdown oocytes, which were still arrested at the Pro-MI/MI stage, even after 10h of culture (60/70). In contrast, the control oocytes had entered anaphase and showed no signals of Bub3 (25/78). To further confirm the phenotype, we analyzed the localization of Mad1, another primary component of SAC in oocytes, and we obtained the same results (Fig. 7B). Detection of Bub3 and Mad1 signals at kinetochores indicates that the SAC (Spindle Assembly Checkpoint) was activated in the CDC6-siRNA injection group.

**Figure 7.**
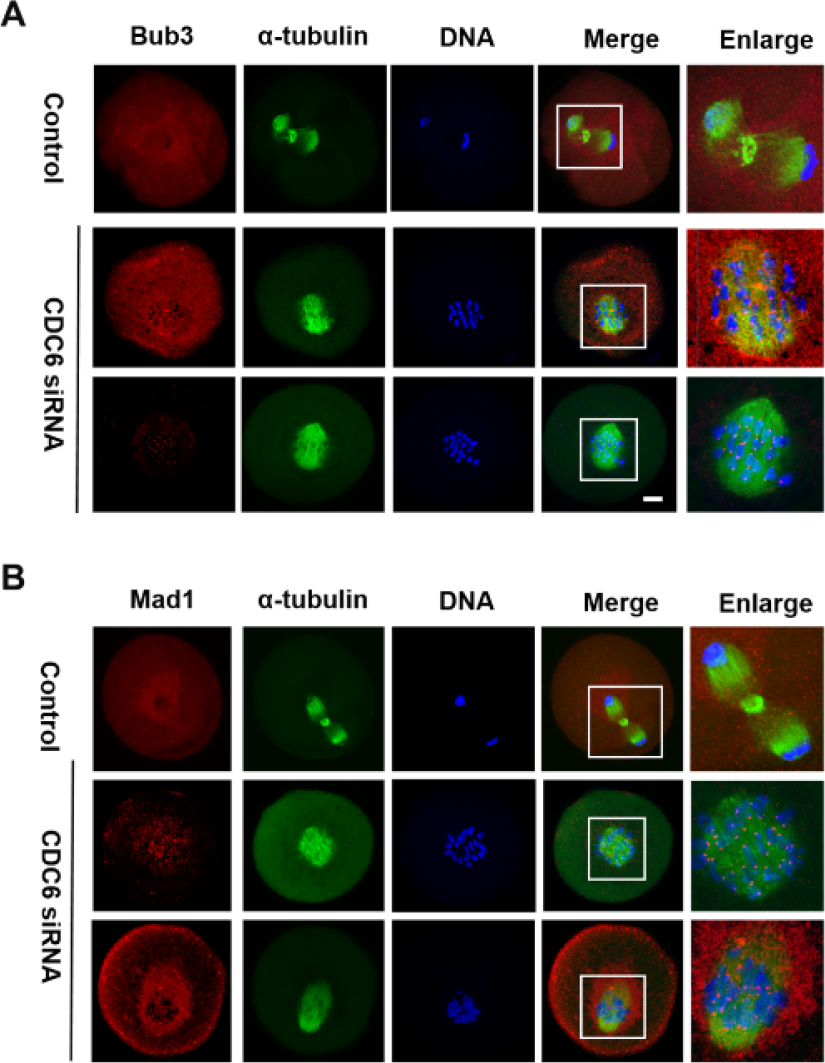
CDC6 knockdown causes activation of SAC. Oocytes from the CDC6 siRNA and control-siRNA injection group were cultured in M2 medium for 10h. (A) Confocal microscopy showing the detection of Bub3 on chromosomes in CDC6-depleted oocytes. Bub3, red; α-tubulin, green; DNA, blue. Bar=20μm. (B) Mad1 fluorescence staining was used to confirm the phenotype. Mad1, red; α-tubulin, green; DNA, blue. Bar=20μm.

### CDC6 knockdown perturbs the metaphase-anaphase transition of mouse oocytes as revealed by time-lapse imaging

Live-cell imaging further showed that in the control group, a clear bipolar spindle was visible at about 4h following GVBD, and it slowly migrated toward the oocyte cortex, then a clear anaphase/telophase stage was observed, followed by rapid polar body extrusion at about 9h following GVBD (Fig. 8A; Movie 1). In contrast, in the CDC6-siRNA injection group, morphologically defective spindles were observed and chromosomes failed to align at the middle plate of the spindle, and thus failed to separate. Oocytes remained at the Pro-MI/MI stage even at 12h following GVBD. Additionally, we could not observe first polar body extrusion (Fig 8B; Movie 2).

**Figure 8.**
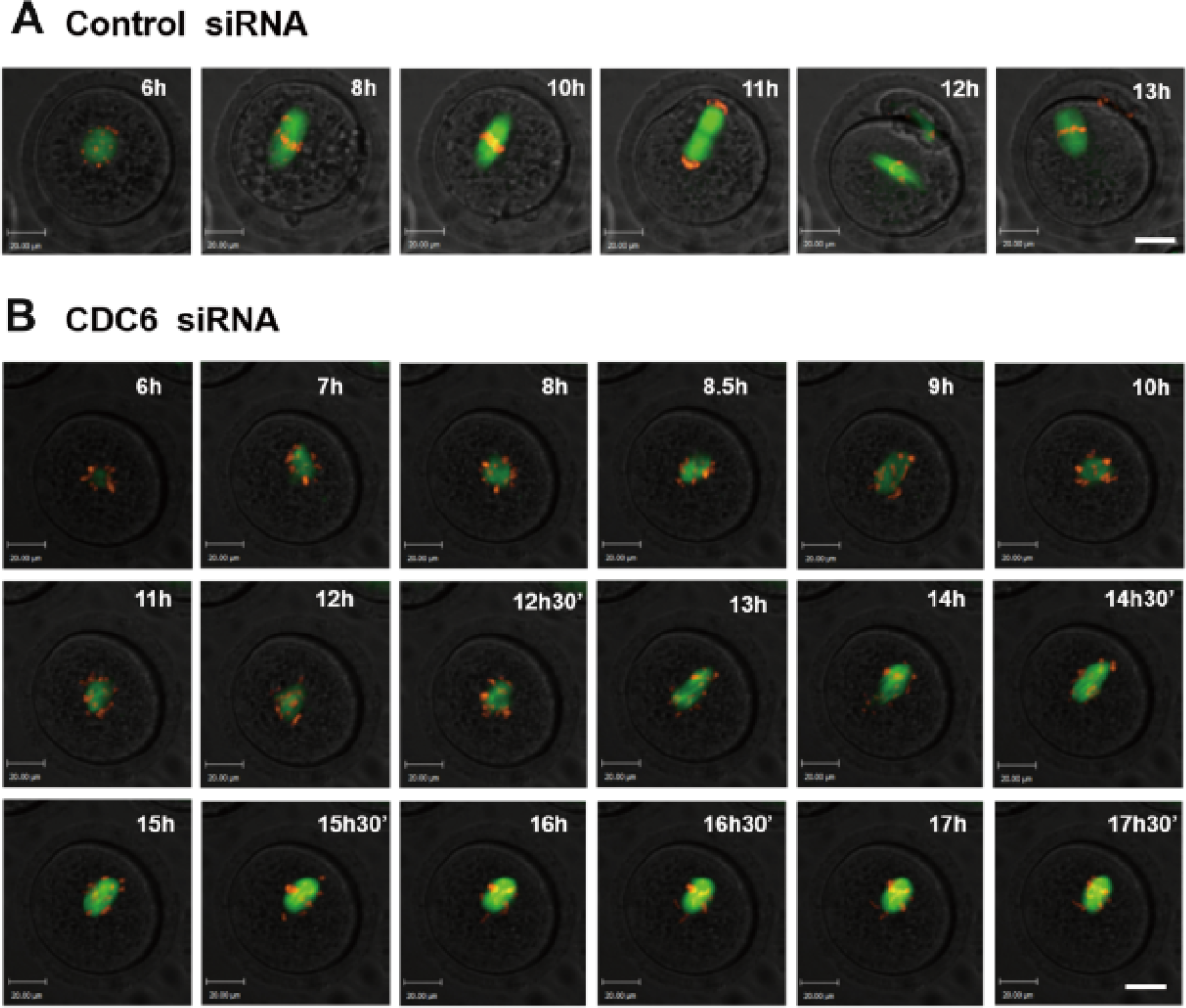
Depletion of CDC6 causes abnormal spindle and misaligned chromosomes, and disturbs the metaphase -anaphase transition of mouse oocytes as revealed by time-lapse live cell imaging. **(A)** Oocytes co-injected with β5-tubulin-GFP mRNA, H2B-RFP mRNA and control siRNA. Spindle (fluorescent tubulin) and DNA (fluorescent H2B) images in a typical control oocyte during in vitro maturation. Time points indicate the time-lapse from about 3-4h after GVBD. (B) Oocytes were co-injected with β5-tubulin-GFP mRNA, H2B-RFP mRNA and CDC6 siRNA. Shown are images of oocytes with abnormal spindles, misaligned chromosomes, unsuccessful chromatin separation and failure of PB1 extrusion. α-tubulin, green; DNA, red. Bar=20μm.

## Discussion

Two critical steps of oocyte meiotic maturation are the G2/M transition (GVBD) and metaphase-to-anaphase transition of first meiosis. In the present study, CDC6, an essential regulator of DNA replication initiation in eukaryotic cells (Diffley, 2004), was shown to regulate these two major events of oocyte meiotic maturation. We firstly show that loss of CDC6 facilitates GVBD in mouse oocytes, which is involved in the regulation of Cdh1 and cyclinB1 expression and thus CDC2 activity. Furthermore, depletion of CDC6 causes abnormal MTOC protein localization with disrupted meiotic spindles and misaligned chromosomes, which causes SAC activation and Pro-MI/MI stage arrest.

It has been shown that CDC6 is located in the nucleus and is relocalized to the cytoplasm when cyclin A/CDK2 is activated in the G1 phase (Petersen et al., 2014). CDC6 was reported to be expressed stably in prophase and being degraded by proteasome activity shortly before entry into meiosis I in Xenopus oocytes (Daldello et al., 2015). While CDC6 and Plk1 colocalized at the spindle pole in metaphase and at the central spindle in anaphase in mitosis (Hyungshin and Erikson, 2010). Here, we showed that CDC6 was expressed at a similar level at the GV, GVBD and MI stages, with a decrease at the MII stage in mouse oocytes. We found that CDC6 was concentrated in the germinal vesicle and distributed along the entire meiotic spindle from Pro-MI to MII stages, suggesting that CDC6 might have different functions in the meiotic events of mouse oocytes.

Overexpression of CDC6 during the late S phase prevents cyclin B/CDK1 activation and leads to cell-cycle arrest at the G2/M–phase transition in mitosis of yeast and mammalian cells(Lorena et al., 2014). In mouse oocytes, it was shown that resumption of meiosis is not arrested by overexpression of CDC6 (Anger and Stein, 2005). Here, we knocked down the expression of CDC6 in GV stage oocytes by siRNA microinjection. Depletion of CDC6 protein caused upregulation of the resumption of meiotic competency, and the ability of oocytes in the CDC6-depletion group to remain in a prophase I arrest in vitro was compromised, despite the addition of milrinone in the culture medium (Fig.2 C). This result indicates that CDC6 activity is essential for maintaining oocyte meiotic quiescence.

It is well known that MPF is an important regulator of oocyte meiotic resumption (Jones, 2004). MPF is a heterodimer composed of a catalytic CDK1 and a regulatory cyclin B1 subunit (Jones, 2004). Increase of cyclin B1 at the GV stage can activate CDK1, and override prophase I arrest. APC/C^Cdh1^ is essential for maintaining G2 arrest through destruction of cyclin B1 (Reis et al., 2007). Milirinone is able to raise the concentration of cyclic adenosine monophosphate (cAMP) and inhibit CDK1 phosphorylation (Norris et al., 2009). It is important to note that although meiosis and G2 arrest is often thought to be primarily due to inhibitory CDK1 phosphorylation, APC^Cdh1^ inhibition or overexpression of cyclin B1 can override any cAMP-imposed arrest (Jones, 2004). Inhibition of CDK1, characterized by phosphorylation of the Tyr15 of CDK1 during G2 is achieved through activation of CDK1-inhibiting kinases (Myt1/Wee1) and suppression of CDK1-activating phosphatases (Han and Conti, 2006). Consequently, alterations in cyclin B1 level and CDK1 activity characterize many conditions that influence meiotic resumption (Holt et al., 2011).

Recently, it has been reported that APC/C^Cdh1^ activity is required to decrease cyclin B1 levels in oocytes during prophase I arrest (Holt et al., 2011). Similarly, APC^Cdh1^ activity in oocytes has been shown to be reduced by Emil and Securin that can inhibit cyclin B1 degradation, or can be up-regulated by Cdc14b, to hasten cyclin B1 degradation (Petros et al., 2008; Marangos and Carroll, 2007). Furthermore, in budding yeast, CDC6 and Cdh1 can cooperate to regulate clb-cdc28 activity and the effect of loss in their activities may result in a failure of cytosis (Archambault et al., 2003).The double mutant Cdh1^loxp/loxp^ and Swis ^loxp/loxp^ showed a reduction in viability, which is also rescued by moderate CDC6 overexpression (Archambault et al., 2003).

Our results show elevation of cyclin B1 levels and activity of CDC2 following CDC6 depletion (Fig.3D). Furthermore, we show that depletion of CDC6 decreases the level of Cdh1, which may be the cause of the increase in cyclin B1 levels. Importantly, accelerated meiotic resumption induced by CDC6 knockdown could be overcome by depletion of cyclin B1 (Fig. 3C). Our results indicate that the CDC6-dependent regulation of CDK1 /cyclin B1 in mouse oocytes is important for the control of meiotic resumption.

These results were further supported by live cell imaging following injection of exogenous cyclin B1-GFP mRNA into the oocytes being arrested in 0.75μM milrinone (which was the minimum concentration to maintain meiotic arrest in mouse oocytes). In the live imaging, the cyclin B1-GFP accumulation is increased significantly in the CDC6 depletion group compared to the control group (Fig4 A,B). While, we found no significant change in cyclin B1 mRNA from real time PCR. Furthermore, raising nuclear cyclin B1 concentrations before GVBD, was a mechanism of inducing the G2/M transition (Holt et al., 2010). Additionally, APC^Cdh1^ activity appeared higher in the nucleus and it guards against cyclin B1 nuclear accumulation (Holt et al., 2010). The translocation of cyclin B1-GFP into the nucleus was increased in the CDC6-depleted oocytes, despite the addition of milrinone, indicative of enhanced nuclear entry, which explains the accelerated meiotic resumption, together with a decreased level of Cdh1 (Fig4. B, C). Our results show that CDC6 is required for G2 arrest in oocytes and that it is indispensable for repressing cyclin B1 levels and nuclear cyclin B1 concentration, which in turn plays an important role in the MPF activation.

A recent study showed that CDC6 inhibits the microtubule-organizing activity of the centrosome in mitosis (Lee et al., 2017). In Hela cells, phosphorylation of CDC6 by Plk1 is required for CDK1 and cyclin B inhibition, which is necessary for separase activation and exit from mitosis (Hyungshin and Erikson, 2010). During the MI-MII transition, there is a feedback loop between CDC6 and CDK1/cyclin B complex that cooperate;CDC6 inhibits CDK1/cyclinB activity, whereas accumulation of B-type cyclins stabilizes CDC6 (Daldello et al., 2015). Recent reports show that mitotic CDC6 stabilize APC/C substrates and affects modulation of APC/C ^cdc20^ in anaphase (Boronat and Campbell, 2007). In this study, we found that knockdown of CDC6 causes spindle assembly defects and extensive chromosome misalignment with predominant spindle defects including mini-spindles and disorganized spindle poles. These results suggest that CDC6 may function in spindle assembly in meiosis. The detailed mechanism remains far from understood. In mouse oocytes, which lack centrioles, the formation of meiotic spindles relies on MTOCs that are localized at the spindle poles (Christiane and Yixian, 2006). MTOCs distributed along the entire spindle at the pro-MI stage before clustering toward the two poles at the MI stages (Manuel et al., 2010). γ-tubulin is a major component of MTOCs and it is important for nucleating microtubules (Sobel and Snyder, 1995). In the present study, we found that CDC6 depletion caused abnormal spindles and chromosome misalignment. Depletion of CDC6 resulted in displacement of γ-tubulin from spindle poles, implying that CDC6 may contribute to spindle organization by partially impairing the function of MTOCs.

Furthermore, CDC6 knockdown prevented the metaphase-to-anaphase transition and chromosome segregation of the first meiosis. Cyclin B1 and CDK1 play important roles in regulating the metaphase-to-anaphase transition in meiosis (Herbert et al., 2003). Once cyclin B1 is degraded by APC/C^Cdc20^, CDK1 activity declines rapidly, leading to exit from the M phase (Herbert et al., 2003). In yeast, the mitotic CDC6 stabilizes APC/C^Cdc20^ substrates and affects modulation of APC/C^Cdc20^ (Boronat and Campbell, 2007). In Hela cells, CDC6 interacts with CDK1 and cyclin B, which is necessary for separase activation and exit from mitosis (Hyungshin and Erikson, 2010). Here, we found that oocytes in the CDC6 depletion group displayed a higher expression of cyclin B1 and higher CDK1 activity than oocytes in the control group at 10h of culture, which indicates that the APC/C^Cdc20^ may be inhibited (Fig. 6 E,F).

SAC proteins such as mitotic arrest-deficient-1 (Mad1), Mad2, budding uninhibited by benzimidazole-I (Bub1), Bub3 and BubR1 have been shown to prevent anaphase onset until all kinetochores of chromosomes are properly attached to the spindle (Li and Murray, 1991, Hoyt et al., 1991). SAC proteins also play important roles in oocyte meiotic maturation (Althoff et al., 2012). Our results show that CDC6 knockdown caused significantly abnormal spindles, which resulted in improper attachments of microtubules to chromosomes. In addition, compared with the control oocytes, CDC6 depleted oocytes were arrested at the Pro-MI/MI stage, and PB1 failed to extrude. These results were further supported by time-lapse microscopy, which revealed that the CDC6-depleted oocytes remained arrested in metaphase and chromosomes failed to separate. Together, these results prompted us to examine whether CDC6 knockdown may activate SAC. Bub3 and Mad1 signals were detected in MI-arrested oocytes in the CDC6 siRNA injection group after 10h of culture, while in the control group, which entered anaphase, the signals were absent. Detection of Bub3 and Mad1 indicated that MI arrest was due to SAC activation caused by CDC6 depletion.

In conclusion, our study shows that CDC6 plays a key role in oocyte GV arrest in the mouse. Additionally, CDC6 cooperates with γ-tubulin to regulate spindle organization, and thus metaphase-to-anaphase transition during the mouse oocyte meiotic maturation process.

## Materials and Methods

### Antibodies and Reagents

Rabbit polyclonal anti-CDC6 antibody was obtained from GeneTex (GTX108979); mouse monoclonal anti-α-tubulin-FITC antibody was obtained from Sigma-Aldrich Co (Cat# F2168); mouse monoclonal anti-γ-tubulin antibody was purchased from Sigma-Aldrich Co (Cat#T6557); rabbit polyclonal anti-bub3 antibody and mouse monoclonal anti-Mad1 antibody were obtained from Santa Cruz Biotechnology (Cat#Sc-28258, Cat#Sc-139025). Mouse monoclonal anti-Cyclin B1 was obtained from Abcam (ab72); mouse monoclonal anti-β-actin antibod was obtained from Santa Cruz (sc-8432); rabbit polyclonal anti-phospho-CDC2(CDK1)-Tyr15 phosphorylated antibody was obtained from Abclonal; rabbit polyclonal anti-FZR1 antibody was obtained from Abcam (ab17038). Alexa Fluor@ 488-conjugate Goat anti-Rabbit IgG (H+ L) and Alexa Fluor @594-conjugate Goat anti-Rabbit IgG (H+ L) were produced by Termo Fisher Scientific (Catalog# A-11008, Catalog# A-11012); TRITC-conjugated goat anti-mouse IgG (H*L) was produced by Jackson ImmunoResearch Laboratories, Inc. and subpackaged by Zhongshan Golden Bridge Biotechnology Co. LTD (Cat#Zf-0313).

All other reagents were purchased from Sigma Aldrich except for those specifically mentioned.

### Ethics statement

Care and handling of 6-8 week-old ICR mice was conducted in accordance with the Animal Research Committee of the Institute of Zoology (IOZ), Chinese Academy of Sciences. ICR mice were housed in an animal care facility which holds a license from the experimental animal committee of the city of Beijing. The mice were sacrificed by cervical dislocation and the only procedure performed on the sacrificed animals was the collection of oocytes from their ovaries.

### Oocyte collection and culture

Immature oocytes with intact germinal vesicle (GV) were collected in M2 medium (Sigma) supplemented with or without 200μM IBMX. IBMX was used to maintain oocytes at the GV stage. The oocytes were placed in M2 medium under liquid paraffin oil at 37□ in an atmosphere of 5% CO_2_ in air. At different times of further culture, oocytes were collected for drug treatment, microinjection, Western blotting and immunofluorescence.

### Real-time PCR

Total RNA was extracted from 100 oocytes with RNeasy micro purification kit (Qiagen). The first strand cDNA was generated with M-MLV first strand cDNA synthesis kit (Takara), using oligo (dT) primers. The primers used to amplify the CDC6 fragment are listed as follows: Forward: 5’-GACACAAGCTACCATCGGTTT-3’; Reverse: 5’-CAGGCTGGACGTTTCTAAGTTT-3’. GAPDH was chosen as a reference gene, and the primers were: 5’-CCCCAATGTGTCCGTCGTG-3’; Reverse:5’-TGCCTGCTTCACCACCTTCT-3’. We used SYBR Premix (Kangwei) in Roche Light Cycler 480.The program for real-time PCR was 95□ for 5 seconds and 60□ for 30 seconds. Analysis of relative gene expression was measured by the real-time quantitative PCR and the 2C-Delta Delta C (T)) method.

### Immunofluorescence and confocal microscopy

Oocytes were fixed with 4% paraformaldehyde in PBS buffer (pH 7.4) at room temperature for 30min and permeabilized with 0.5% Triton X-100 for 20 min. After being blocked in 1% BSA for 1h at room temperature, the oocytes were then incubated overnight at 4□ with rabbit polyclonal anti-CDC6 antibody (Genetex, 1:50), mouse polyclonal anti-α-tubulin-FITC (Sigma, 1:100), mouse anti-γ-tubulin antibody (Sigma, 1:100), rabbit polyclonal anti-Bub3 antibody (Santa Cruz Biotechnology, 1:50) and mouse monoclonal anti-Mad1 antibody (Santa Cruz, 1:50), respectively..

After three washes in washing buffer (0.1% Tween 20 and 0.01% Triton X-100 in PBS), the oocytes were then labeled with F488-conjugated goat-anti-rabbit IgG (1:100), F594-conjugated goat-anti-rabbit IgG (1:100) or Tritc-conjugated goat anti-mouse IgG (1:100) for 2h at room temperature. Finally, the oocytes were stained with Hoechst 33342 for 15min to detect DNA. Then, the oocytes were mounted on glass slides with anti-fade mounting medium (DABCO) to retard photobleaching and visualized with Cal Zeiss LSM 780 confocal microscope.

### Immunoblotting analysis

Samples (each containing 200 oocytes) were collected in 2×SDS loading buffer and boiled for 5 min. The proteins were separated by SDS-PAGE and then transferred to polyvinylidene fluoride (PVDF) membranes. Thereafter, the membranes were blocked in TBST containing 5% bovine serum albumin for 1h at room temperature, followed by incubation overnight at 4□ with Rabbit polyclonal anti-CDC6 (1:1000), mouse monoclonal anti-cyclin B1 (1:1000), rabbit polyclonal anti-phospho-Cdc2 (CDK1)-Tyr15 (1:1000), rabbit polyclonal anti-FZR1 antibody (1:1000) or rabbit monoclonal anti-β-actin antibody (1:2000). After three washes with TBST, 10 minutes each, the membranes were incubated with horseradish peroxidase (HFP)-conjugated goat anti-rabbit IgG (1:1000) and HRP-conjugated goat anti-mouse IgG (1:1000), respectively, for 1 hour at 37□. Finally, the membranes were washed three times in TBST and processed with the enhanced chemiluminescence (ECL)-detection system (Amersham, Piscataway, NJ).

### Chromosome spreading

The oocytes were placed in acid Tyrode’s solution (Sigma) for 1 minute at 37□ to remove the zona pellucida. After a brief recovery in M2 medium, the oocytes were placed onto glass slides and fixed in a solution of 1% paraformaldehyde in distilled H_2_O (PH 9.2) containing 0.15% Triton X-100 and 3mM dithiothreitol. Chromosomes on the slides were stained with 10µg/ml Hoechst 33342 and the specimens were mounted for immunofluorescence microscopy observation.

### Microinjection of mRNA and siRNA

Microinjections were performed using a Nikon Diaphot ECLIPSE TE 300 (Nikon UK Ltd), and finished within 30 min. A volume of 10μM CDC6 siRNA (Santa Cruz Biotechnology, SC-3504C) or cyclin B1 siRNA (Genepharma, China) was microinjection into the cytoplasm of oocytes to deplete CDC6 or cyclin B1. The same amount of scrambled siRNA was injected as control.

To examine how CDC6 depletion perturbed the meiotic division, we co-injected β-tubulin-GFP mRNA and H2B-RFP mRNA which were synthesized according to references Verlhac et al., 2000 and Wang et al., 2008, with 20μM CDC6 siRNA or control siRNA into GV oocytes as described above (Verlhac et al., 2000, Wang et al., 2008). After injection, the oocytes were arrested at the GV stage in M2 medium containing 200μm IBMX for 24h to allow depletion of CDC6. Then the oocytes were thoroughly washed and transferred to IBMX-free medium. To examine cyclin B1-GFP dynamics, 20ng/μl cyclin B1 mRNA was injected into GV oocytes.

Each oocyte was microinjected with approximately 10pl CDC6 siRNA, control siRNA, β5-tubulin-GFP mRNA and H2B-RFP mRNA with CDC6 siRNA or β5-tubulin-GFP mRNA and H2B-RFP mRNA with control siRNA. Each experiment contained three separate replicate groups and approximately 300 oocytes were injected in each group.

### Time-lapse live-imaging experiments

Cyclin B1-GFP or microtubule and chromosome dynamics were filmed on a Perkin Elmer precisely Ultra VIEW VOX Confocal Imaging System. A narrow band passed EGFP and RFP filter sets and a 30% cut neutral density filter from chroma were used. Exposure time was set ranging between 300-8000ms depending on cyclin B1-GFP, tubulin-GFP and DNA-RFP fluorescence levels. The acquisition of digital time-lapse images was controlled by IP lab (Scanalytics) or AQM6 (Andor/Kinetic-imaging) software packages. Confocal images of spindles and chromosomes in live oocytes were obtained with a 20/10 oil objective on a spinning disk confocal microscope (Perkin Elmer).

### Statistical analysis

For all experiments, at least three replicates were performed. Data are expressed as means± SEM, and the number of oocytes observed are given in parentheses as (n=). Statistical analyses were conducted by Student’s t-test with SPSS 13.0 software (SPSS Inc.). Fluorescence intensity statistics were conducted using the Image J (NIH). P<0.05 was considered statistically significant.

## Acknowledgements

This work was supported by the National Natural Science Foundation of China [31530049]; The Research Team of Female Reproductive Health and Fertility Preservation [SZSM201612065]; Project for Medical Discipline Advancement of Health and Family Planning commission of Shenzhen Municipality [SZXJ2017003]; and Natural Science Foundation of Guangdong Province [2018A030310673].

## Author contributions

Zi-Yun Yi designed and conceived the experiments; Zi-Yun Yi wrote and all authors reviewed the manuscript. Wei-Ping Qian, Qing-Yuan Sun, Chun-Hui Zhang, and Jie Qiao provided professional advice on experimental design and paper writing. Heide Schatten edited the manuscript. Tie-Gang Meng, Xue-Shan Ma, Jian Li, Yi Hou, Ying-Chun Ouyang provided assistance with experiment performance.

---

**Figure S1.**
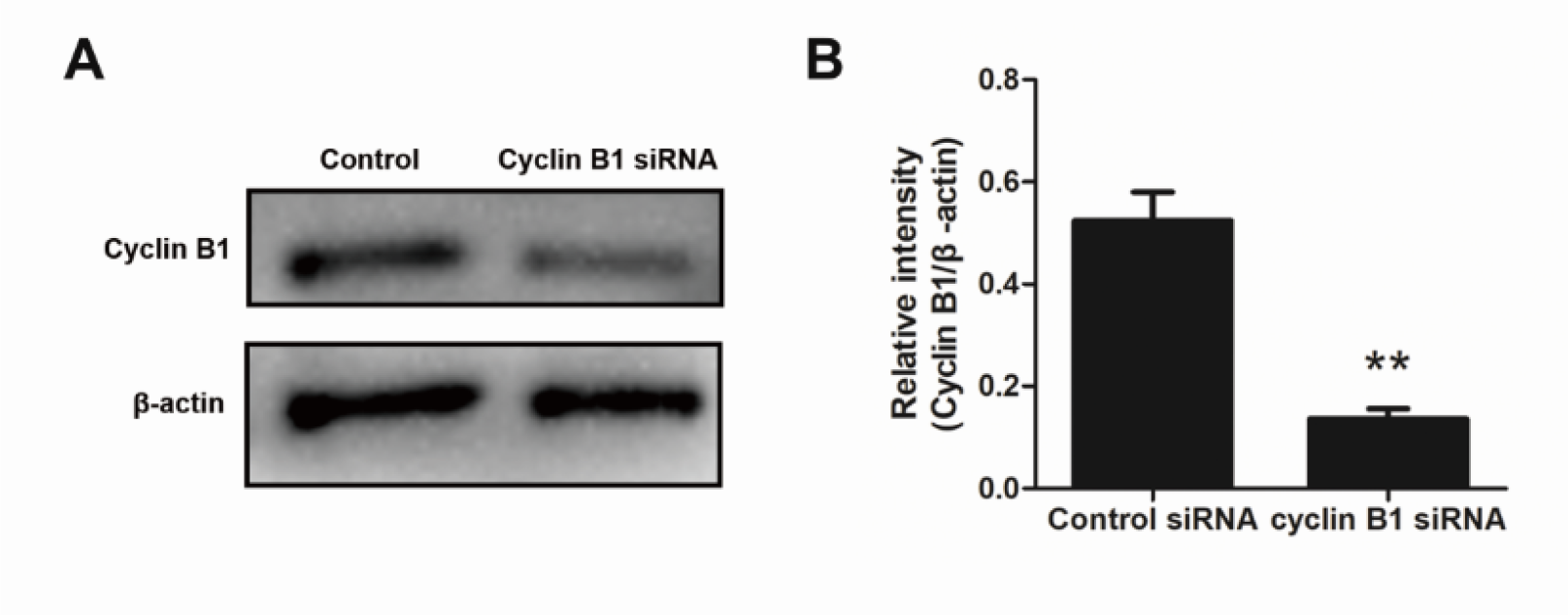
Western blot of cyclin B1 in the oocytes microinjected with cyclin B1 siRNA or control siRNA. The oocytes were arrested in M2 medium containing 200μM IBMX for 24h. The molecular mass of cyclin B1 is 55kDa and that of β-actin is 43kDa. A total of 200 oocytes were used in each lane. The relative intensity of cyclin B1 was assessed by gray-scale analysis using the software Quantity One (Bio-Rad). Levels of staining intensity were normalized to levels of β-actin. Data are presented as mean±SEM of 3 independent experiments (**,p<0.01).

